# Intrathecal infusion of hypertonic fluid enables CSF Flow Enhancement (CFE) to facilitate nanoparticle delivery to the brain and spinal cord

**DOI:** 10.64898/2026.06.29.735409

**Authors:** Oluwatobi Babayemi, Kha-Uyen Dam, Chung-Fan Kuo, Olivia Mihalek, Elena A. Andreyko, Constance J Mietus, Shaokuan Zheng, Hong Wei Yang, Rachael W. Sirianni

## Abstract

Intrathecal (IT) drug delivery, i.e., the infusion of substances directly into cerebrospinal fluid (CSF) by lumbar, ventricular, or cisternal access points, is one method that can be used to bypass the blood brain barrier (BBB), however, IT-administered substances also suffer from rapid turnover and poor tissue penetration. Although nanoparticles and colloids can circulate within the subarachnoid space to sustain the levels of encapsulated drug in CSF, their access to deep tissue regions remains incomplete. Here, we present a new method for enhancing CNS delivery of IT-administered nanoparticles. CSF Flow Enhancement (CFE) refers to the manipulation of CSF production, distribution, and clearance for therapeutic purposes. We tested the overarching hypothesis that infusion of hypertonic fluid adjacent to the choroid plexus would enhance fluid production and movement to improve the CNS delivery of IT-administered nanoparticles. Model polystyrene nanoparticles (100nm) were solubilized in aCSF of increasing tonicity (1-9X tonicity) and infused into the cisterna magna, after which tissues were removed to examine delivery to CNS tissues and peripheral organs. Our results demonstrate that an infusion of up to 4X hypertonic aCSF in 10uL is well tolerated and yields significant improvements in CNS localization of co-administered nanoparticles, more than doubling the delivery of nanoparticles to the ventral surfaces of the brain and sometimes dramatic (up to 10-fold) increases in delivery to specific tissue regions and surfaces of the CNS. Significantly, we provide early evidence that modulation of tonicity can define the parenchymal fate of IT administered colloids: while nanoparticles were not detected in the brain parenchyma of mice that received a standard infusion, parenchymal delivery was observed for the 2X condition, and extensive perivascular infiltration of nanoparticles was observed for the 4X condition. Lastly, we show that the delivery improvements achieved by CFE are generalizable across multiple sizes of polystyrene nanoparticle (20, 40, or 100nm). Collectively, this work describes a tonicity-based approach for achieving CFE by the intrathecal route, which we posit is a useful and potentially generalizable approach for improving CNS drug delivery.

## 1. Introduction

Intrathecal (IT) administration refers to the infusion of substances directly into cerebrospinal fluid (CSF), and is a method of central nervous system (CNS) drug delivery that bypasses the blood-brain barrier (BBB) and blood-spinal cord barrier (BSCB)^1^. While this approach can be used to improve locoregional drug retention in the CNS for some lipophilic agents, particularly for pain management and chemotherapy, conventional IT drug delivery is limited by poor parenchymal penetration and high rates of drug clearance due to rapid turnover of CSF^2^. Encapsulation of bioactive molecules within nanoparticles (NPs) is one potential solution. Because NPs circulate readily but clear slowly from the subarachnoid space^2, 3^, use of a nanoparticle delivery system can sustain drug levels within CSF or tissues of the CNS after IT administration. However, our prior work shows that although intrathecally administered NPs distribute rapidly within the subarachnoid space and can reach some parenchymal tissues, the total quantity of NPs that reach deep tissue targets within the brain is low^2–4^. To improve drug delivery by the intrathecal route, new strategies will be needed.

This work addresses an emerging approach for CNS drug delivery termed CSF Flow Enhancement, or CFE^5^. CFE refers to manipulations that alter production, flow, or clearance of CSF. A multitude of approaches have been used to manipulate CSF, including pharmacological agents, focused ultrasound, and even exercise or repeated flexion of the neck. Importantly, growing evidence shows that manipulation of CNS hydrodynamics via CFE may be useful for therapeutic purposes. For example, a longstanding approach is to administer systemic hypertonic saline or mannitol, which can enhance permeability of the BBB to enable systemically circulating agents to enter the parenchyma but that also facilitates fluid egress from the CNS^6^. Clinically, this strategy is used to treat high intracranial pressure (ICP); by sequestering fluid from brain parenchyma into the blood, ICP is lowered, decreasing brain edema and swelling^7, 8^. In one recent report, intrathecal mannitol was supplied intrathecally instead of intravenously, which enhanced antibody penetration along perivascular channels. We have shown in prior work that recirculation of CSF can improve the distribution of methotrexate in the CNS after intrathecal administration^5^. These and other examples provide support for the concept that the distribution of CSF can be manipulated to control the distribution of substances within the CNS.

Here, we developed a new method of CFE, focusing specifically on the chemosensory capability of the choroid plexus (CP)^9^. CP is an evolutionarily conserved tissue structure that consists of cuboidal epithelial cells fueled by fenestrated vasculature that lacks a BBB. Importantly, CP tissues possess a remarkable ability to respond rapidly to changes in CSF concentration (including ion concentration, pH, and total osmolality), secreting fresh CSF that turns over to maintain CNS homeostasis^10^. Given growing knowledge that CSF clearance is driven through perivascular routes, i.e., the glymphatic system^11, 12^, we hypothesized that an osmotically triggered increase in CSF production could be used to drive intrathecally administered NPs into deeper tissue targets via perivascular clearance routes. To test this hypothesis, we co-administered hypertonic aCSF (1X-9X native osmolality) with 100nm, fluorescent polystyrene NPs to healthy mice via the intrathecal cisterna magna (IT-CM) route. Using a combination of stereoscopic, epifluorescent, and confocal imaging, we quantified NP delivery across and into CNS tissues. Our results establish tolerable limits of safety for achieving intrathecal CFE via osmotic mechanisms and also demonstrate that CFE can yield major improvements in the tissue and cellular level fate of NPs within the CNS, achieving up to 5-fold higher levels of NP delivery to specific regions of the CNS. Taken collectively, we describe a new, easy to implement approach for enhancing brain and spinal cord delivery of IT-administered colloids.

## 2. Materials and Methods

### 2.1. Materials

NaH_2_PO_4_•2H_2_O (446222500), NHS (16050122), and 100nm carboxylated FluoSpheres^TM^ (Red, λ_ex_/λem = 580/605) (F8801) were sourced from ThermoFisher Scientific (Waltham, MA, USA). Endotoxin-free, ultra-pure water (TMS-011-A), NaCl (S9888), KCl (P3911), MgCl_2_ (M8266), CaCl_2_•H2O (C3881), and Triton X-100 (T8787) were purchased from Sigma-Aldrich (St. Louis, MO, USA). FITC goat anti-rabbit IgG (ab6717) were purchased from Abcam (Waltham, MA, USA). Rat CD31 monoclonal antibody (MA1-40074) and Alexa Fluor® 647 goat anti-rat IgG (A-21247) were purchased from Invitrogen (Carlsbad, CA, USA). Alexa Fluor® 647 (AF) rat anti-mouse CD31 antibody was sourced from BioLegend® (San Diego, CA, USA). Polyethylene (PE) tubing, with 0.011in inner diameter and 0.024in outer diameter, was sourced from Braintree Scientific (PE10) (Braintree, MA, USA).

### 2.2. aCSF preparation

Artificial CSF (aCSF) was prepared using a combination of established protocols^13–16^. To prepare 100mL of stock 10X aCSF, two solutions (solution A and solution B) were combined in a 1:1 ratio. For solution A, a mixture of 125mM NaCl, 3mM KCl, 1mM MgCl_2_ • 6H_2_O, and 2mM CaCl_2_ • H_2_O was dissolved in 50mL endotoxin-free, ultra-pure water. For solution B, 1.25mM NaH_2_PO_4_ • 2H_2_O was dissolved in 50mL endotoxin-free, ultra-pure water. After combining both solutions, the resulting 10X aCSF stock was filtered using a 0.22µm filter to remove any unwanted aggregates and stored at 4°C for further use. For experiments, 10X aCSF aliquots were diluted in water to varying ratios (1X–9X) and titrated as needed to adjust pH.

### 2.3. Nanoparticle preparation

To prepare NPs, 1mL aliquots of 100nm carboxylated FluoSpheres^TM^ (FNPs, starting concentration at 2wt%) were washed three times with 1X DPBS buffer using 10kDa MWCO Amicon-Ultra Centrifugation filters and a Beckman Coulter AllegraX-30R centrifuge (Indianapolis, IN, USA) at 4255 RCF and 25°C. After washing, 1mL of the FNP concentrate was resuspended in 2mL 1X DPBS buffer. FNPs were solubilized in water and combined with 10x aCSF to achieve the desired tonicity.

### 2.4. IT-CM administration

All *in-vivo* studies were conducted in accordance with UMass Chan Medical School policies and approved by the Institutional Animal Care and Use Committee (IACUC) (Protocol ID: IPROTO202300000008 & PROTO202200109). Healthy, 7-to-9-week-old female B6(Cg)-Tyr^c-2J^/J, i.e., C57Bl/6J or albino C57BL/6J mice (strain #: 000058, RRID: IMSR_JAX:000058) (The Jackson Laboratory, Bar Harbor, ME, USA) were used in all experiments. A percutaneous (non-surgical) method was used to access the cisterna magna in mice, as described by Reijneveld et al. (1999)^17^ and reported by us previously^2, 3^. Briefly, mice were anesthetized with isoflurane (2%) and placed on a cylindrical barrel in a prone position, with their necks draped over the barrel and their noses pointing downward at a 30° to 45° angle. Using the thumb and middle finger, the head was immobilized, and the index finger used to palpate the space between the base of the skull and C1. A 33-gauge Hamilton syringe (adjusted with a stopper to set needle length at 2-3mm) was then used to administer 10µL volumes as a slow bolus (∼30-45s).

### 2.5. Euthanasia and Tissue Fixation

At defined time points, mice were deeply anesthetized and underwent transcardial perfusion with heparinized PBS followed by fixation with 4% paraformaldehyde (PFA). Exsanguinated tissues of interest (the brain, spinal cord, cervical lymph nodes (CLNs), inguinal lymph nodes (ILNs), liver (with gallbladder), kidneys, spleen, and lungs) were post-fixed with 4% PFA for 24-48hrs at 4°C, washed 3 times with 1X DPBS, and transferred to 30% sucrose for storage at 4°C until further use. Brain and spinal cords were carefully dissected to maintain the pia mater and to keep exiting nerve roots and the cauda equina intact.

### 2.6. Fluorescent Stereoscopy

Whole tissues, including intact brain, spinal cord, and lymph nodes, were imaged with a Leica M205 FA fluorescent stereoscope fitted with a K5 camera (Leica Microsystems Inc., Deerfield, IL, USA). Image acquisition settings, including exposure, were maintained across samples; only linear adjustments were applied to the resultant images. Brain images were collected on both the dorsal and ventral surfaces. It was not possible to consistently maintain dorsal-ventral orientation for spinal cord imaging due to mild curling and twisting artifacts introduced by variable mouse orientation upon tissue fixation. Spinal cords were therefore imaged on the “top” and “bottom” surfaces, which were then averaged to report a mean value for the spinal cord. Top and bottom signals were similarly averaged for the lymph nodes. An ROI was carefully drawn on the right, ventral surface of the brain to isolate lateral vessels. Images were exported as TIFF files for additional analysis with Fiji software (ImageJ2 version 2.16.0/1.54r). Total fluorescence was measured in various regions of interest and normalized to the 1X group to examine fold-change in FNP signal. Line plots generated by Fiji are reported in AU; to determine fold-change in signal across the rostral-caudal direction, a smoothing function was applied (20 pixels). Line plots were additionally generated for the medial-lateral direction for spinal cord samples.

### 2.7. Magnetic Resonance Imaging (MRI)

Mice (n=2/group) received 1X, 2X, or 4X FNP infusions via the IT-CM route and were imaged at three time points: pre-injection, immediately post injection (0 hrs p.i.), and two hours later (2hrs p.i). For all groups, a 3D coronal T2-weighted scan of the head was performed. A BioSpec 70/30 USR horizontal bore MR system (Bruker Corporation, Billerica, MA, USA) was used to collect coronal, T2-weighted MRI images. Changes in ventricular volume were assessed with a combination of MATLAB (R2024b) and FIJI. First, MRI images were analyzed in Fiji to generate a 2D image stack of the brain. Each image was cropped to isolate the ventricles, ensuring that extraneous signal from surrounding tissue was removed to improve accuracy in subsequent volume quantification. The processed image stack was then analyzed using an in-house MATLAB script. The ventricle ROI was isolated on the basis of pixel intensity, which was then integrated across the entire image stack to create a 3D rendering of the ventricles and estimate volume in µL.

### 2.8. Whole-body fluorescent imaging

A subset of mice that received IT-CM infusion of FNPs with (2X and 4X) and without (1X) CFE were imaged with an IVIS® SpectrumCT (Perkin Elmer, Waltham, MA, USA). Imaging was initiated as soon as possible after compound administration and proceeded in 5-minute increments for 2 hours. These images were analyzed by Living Image™ software, with ROIs defined to surround the whole head, olfactory bulb, cisterna magna, upper spine, lower spine, tail, or paws. For *ex vivo* analyses, the brain, spinal cord, and peripheral tissues were imaged on both the ventral and dorsal surfaces.

### 2.9. Immunohistochemistry (IHC)

After collection, some brain tissues were embedded in OCT compound and cryopreserved at −80°C. Tissues were first sliced into thick, 1mm coronal sections, embedded and frozen in OCT, and then further sliced into 14µm-thick coronal slices on a cryotome for IHC staining. Samples were rehydrated with 1X PBS and subsequently blocked with 4% NHS and 0.3% TritonX-100 in PBS. Specimens were then treated with a mixture of rabbit monoclonal anti-AQP4 and rat monoclonal anti-CD31 (1:200) and maintained at 4°C overnight. Negative controls were also prepared using 4% NHS. Slides were next treated with a mixture of FITC goat anti-rabbit IgG (1:250) and AF594 goat anti-rat IgG (1:500). DAPI nuclear counterstain was applied, and slides were mounted using Fluoromount G (cat# 0100-01, Southern Biotech, Homewood, Alabama, USA) and coverslipped. Slides were imaged at 160X using the Leica M205 stereoscope, with image acquisition and processing parameters carefully maintained across experimental groups.

### 2.10. Stability Studies

FNP stability studies in 1X aCSF and 1X PBS were conducted as previously described^3^. FNP stability was assessed in 1X, 2X, 4X, 6X, and 8X aCSF following incubation at 37°C for 0hrs, 0.5hrs, 1hr, 2hrs, and 4hrs. At each time point, samples were removed and zeta potential, polydispersity index, and hydrodynamic diameter were measured using a NanoBrook 90Plus Zeta (Brookhaven Instruments, Holtsville, NY, USA).

## 3. Results

In the first series of experiments, mice received 10uL of FNPs suspended in media of varying tonicity (1X, 2X, 4X, 6X, or 8X tonicity relatively to standard aCSF) that was administered as a slow bolus by the IT-CM route; whole brains and spinal cords were extracted 2 hours later for assessment of FNP distribution with fluorescent stereoscopy of intact tissues (**Figure 1A**). This imaging method allows visualization of surface, surface-adjacent, and meningeal-vessel-associated FNPs while neglecting signal contributions from the deep parenchyma, arachnoid mater, dura mater, and skull or vertebral bone marrow. Importantly, for these and all following results, exposure and gain settings were adjusted based on control tissue collected from mice that did not receive an FNP injection to eliminate background, and imaging settings were maintained across experimental groups, i.e., the appearance of fluorescent signal in these images is indicative of FNP presence and not an artifact of tissue autofluorescence.

**Figure 1:**
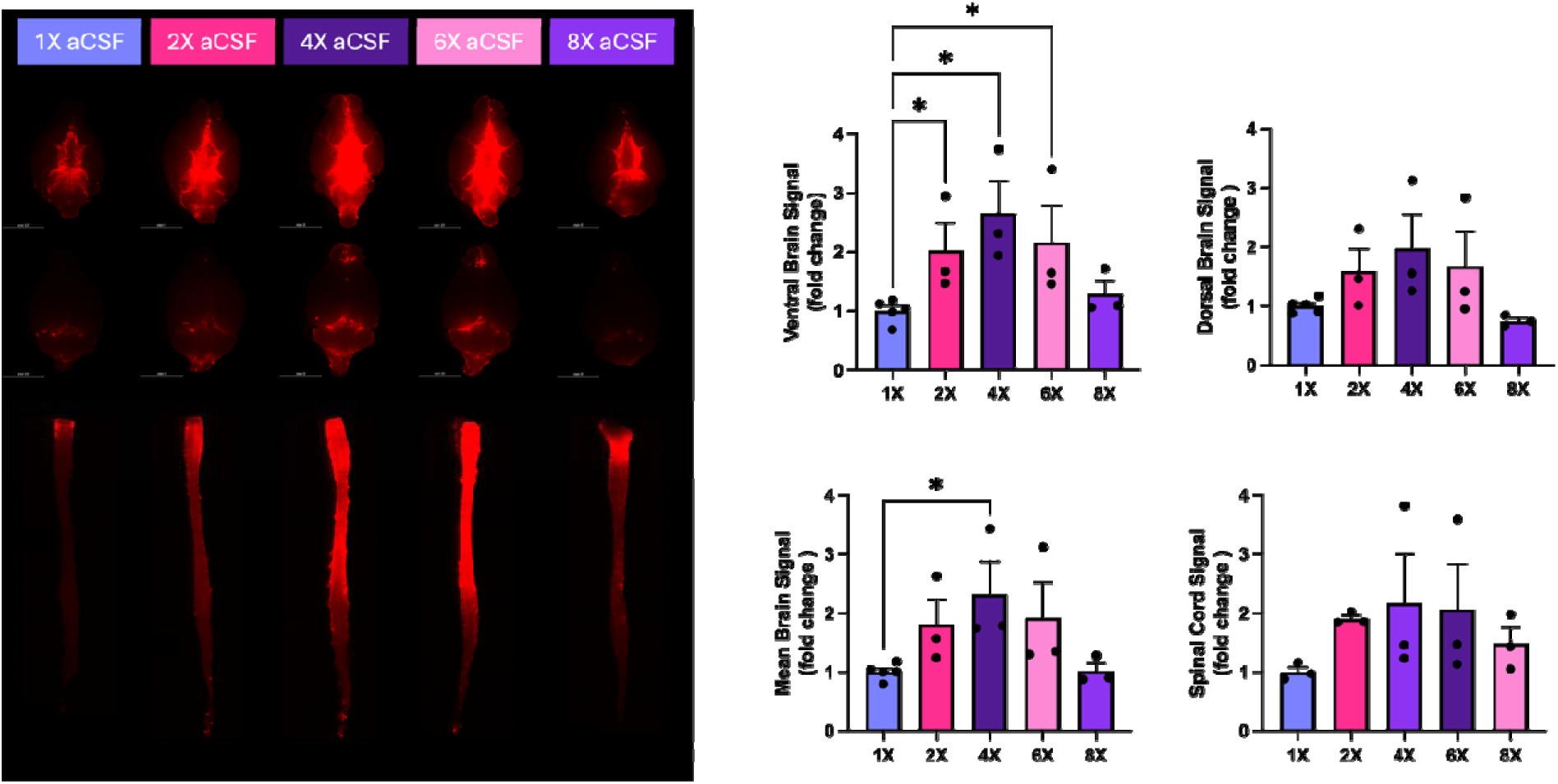
Application of CFE increases FNP association with the surfaces of CNS tissues. Mice (n=3-5 per group) received FNPs diluted in aCSF of varying tonicity (1X, 2X, 4X, 6X, and 8X) as a 10uL infusion by the IT-CM route. When assessed along ventral surfaces, 2X, 4X, and 6X aCSF yielded a statistically significant increase in FNP localization with ventral brain surfaces compared to 1X control. When averaged across both the ventral and dorsal surfaces, 4X aCSF yielded a significant increase in FNP delivery compared to 1X control. There was a tendency for 2X, 4X, and 6X conditions to increase FNP localization with the surfaces of the spinal cord, although these differences were not statistically significant. Data were analyzed by Kruskal-Wallis with Dunn’s test for multiple comparisons. Data show mean plus/minus standard deviation, with significance reported for *p<0.05. Scale bar is 5mm.

As we previously observed^2, 3^, FNPs administered by the IT-CM route in 1X aCSF distributed readily across the neuroaxis, reaching all CSF-exposed surfaces of the brain and spinal cord and localizing in a reproducible manner with the supracerebellar cistern, dense trabeculae networks present on the ventral surface of the brain, the Circle of Willis, and exiting nerve roots of the spinal cord. As the aCSF infusion media tonicity was increased, we observed dose-dependent increases in FNP localization with tissues of the CNS as assessed by signal in each region of interest (ROI, **Figure 1B**). Mice that received 1X infusions exhibited highly reproducible FNP signals, with a subject-to-subject standard deviation of 15.4% for brain delivery (averaging dorsal and ventral views obtained for each mouse); brain delivery was increased by application of CFE conditions compared to 1X control for the 2X, 4X, and 6X groups, reaching 1.8, 2.3, and 1.9 fold-higher FNP signal with associated standard deviations of 70%, 95%, and 98%, respectively. Individual mice exhibited an up to 3 to 6-fold increase in FNP signal for CFE conditions compared to 1X aCSF, and every mouse in the 2X-6X CFE groups exhibited a higher brain signal than each mouse in the 1X aCSF control group. Delivery benefits appeared to reach a nominal maximum in the brain at the 4X condition and in the spinal cord at the 6X condition. There was a near linear correlation between dorsal and ventral signals (**Supplementary Figure 1**), emphasizing that the subject-to-subject variability is likely not a consequence of administration or measurement uncertainty but instead likely represents physiological differences in the extent of CFE achieved in different subjects. Benefits of CFE were apparently maximized at 6X aCSF, with distributions returning close to baseline and a reduction in subject-to-subject variability for the 8X condition.

We next examined the spatial distribution of FNPs along the dorsal and ventral surfaces of CNS tissues (**Figure 2**). Line plots representing the rostral-caudal axis were generated to carefully align major anatomical features of the CNS. Brain line plots were aligned according to the disappearance of signal at the tip of the olfactory bulb and the maximization of signal at the cisterna magna. The spinal cord exhibited unavoidably variability in its length and dorsal-ventral orientation depending on exact positioning during tissue fixation; we therefore aligned spinal cords line plots at the top (cervical region) of the spinal cord to calculate a mean signal representing images collected for both the top and bottom of each sample, and excluded the cauda equina from spatial analyses, as it has highly variable extraction efficiency at dissection.

**Figure 2:**
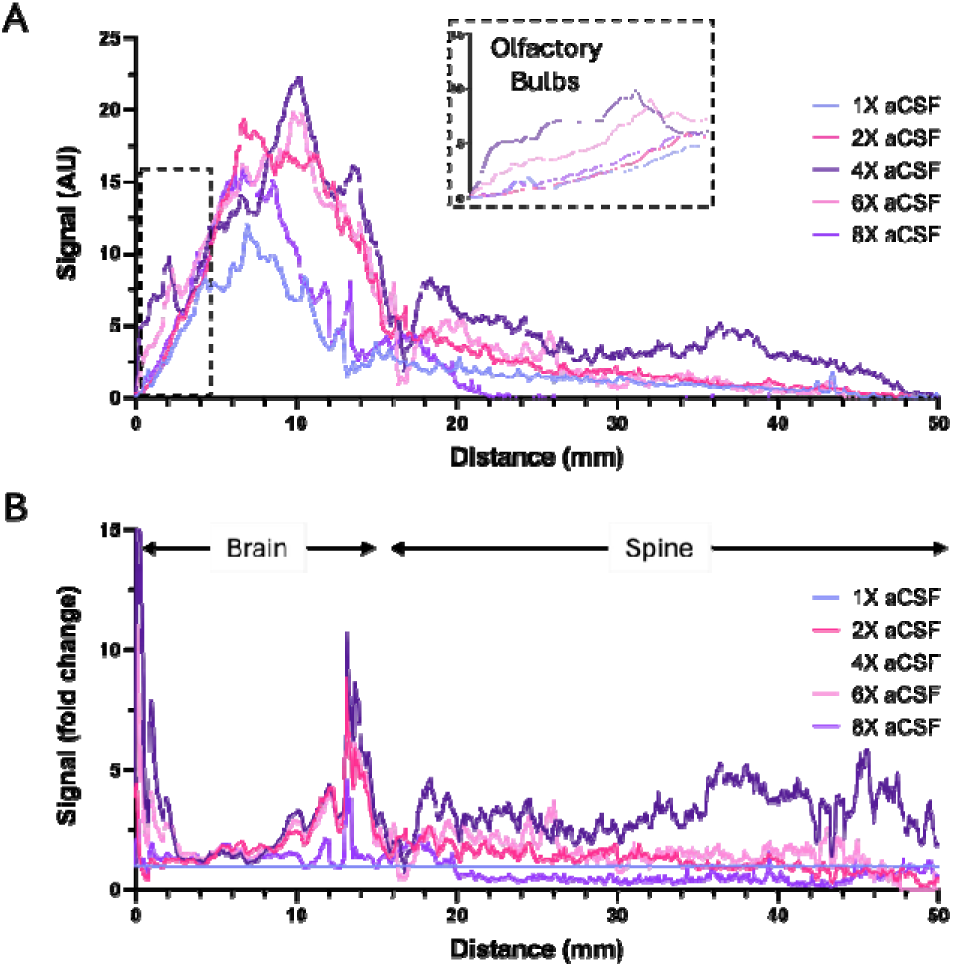
Application of CFE increases the spatial distribution of FNPs across CNS tissues. Mice received FNPs diluted in aCSF of varying tonicity (1X, 2X, 4X, 6X, and 8X) as a 10uL infusion administered by the IT-CM route. Line plots were generated to examine FNP signals across the complete neuroaxis, averaging dorsal and ventral surfaces. (A) When assessed on an absolute scale, CFE conditions consistently increased FNP delivery to both the brain and spinal cord. Particular improvements were seen in the olfactory bulb (shown in inset). (B) Following normalization to the 1X condition, improvements in delivery are consistently observed for 2X, 4X, and 6X conditions, with consistent enhancement observed in the brain and upper spinal cord. Data for both panels show the mean of n=3 mice per group.

Quantitative analyses confirmed that CFE generated differences in FNP spatial distribution that followed similar trends as what was observed for whole-tissue imaging, with higher tonicity of aCSF yielding a higher FNP signal associated with the surfaces of the brain and spinal cord (**Figure 2A**); the impact of CFE is further evident when smoothed plot profiles are expressed as a fold change to the mean of the 1X condition, emphasizing increased delivery through the brain and spinal cord with region. Regions of interest that were particularly impacted by CFE included the olfactory bulbs, the region anterior to the cisterna magna. The cisterna magna contains signal contributions from the Circle of Willis and supracerebellar signals, which is also where 2X, 4X, and 6X conditions also yield high benefit over 1X and 8X.

We next examined potential CNS outflow pathways, extracting 5-7 CLNs and 2 ILNs from each subject (**Figure 3**). This analysis yielded tonicity-dependent differences in FNP egress from the CNS, with 2X, 4X, and 6X CFE conditions yielding relatively higher CLN delivery (2.1±0.65, 2.7±1.1, and 2.0±1.6, respectively; fold-difference relative to control) but no differences in ILN delivery as a function of infusion media. As before, we observed a relatively high level of intersubject variability; the subjects with the highest CLN delivery were also those with the highest CNS tissue delivery, and there was no relationship between CLN and ILN delivery on a per-subject basis (see **Supplementary Figure 2**).

**Figure 3:**
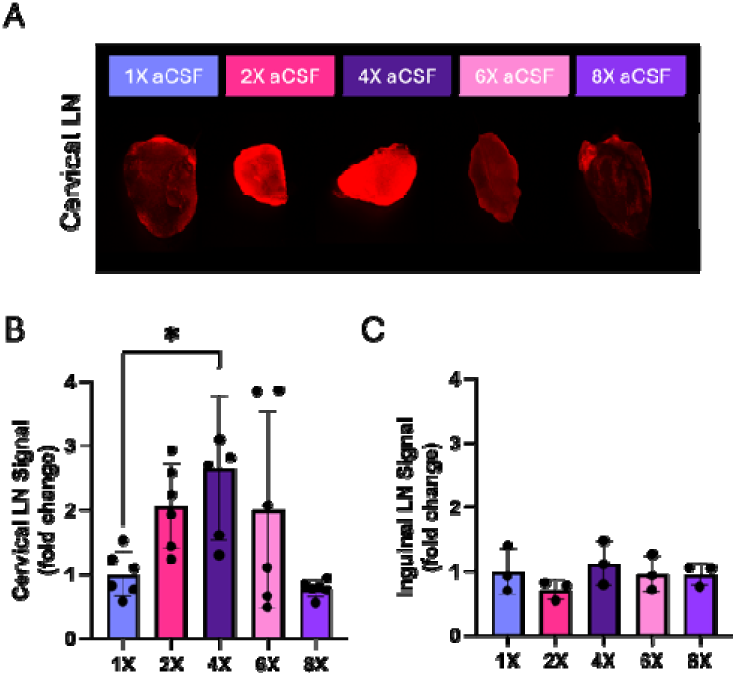
Application of CFE increases FNP localization with cervical but not inguinal lymph nodes. Mice received FNPs diluted in aCSF of varying tonicity (1X, 2X, 4X, 6X, and 8X) as a 10uL infusion administered by the IT-CM route. Lymph nodes were extracted and imaged for whole-tissue fluorescence. (A) Representative images are shown for the cervical lymph nodes. (B) Quantification of total FNP signal yields a significant increase in CLN localization for the 4X group compared to 1X conditions; left/right CLNs are aggregated for a total of 6 samples from 3 mice. (C) There was no difference in delivery to inguinal lymph nodes as a function of infusion medium tonicity. Data were analyzed by Kruskal-Wallis with Dunn’s test for multiple comparisons. Data show mean plus/minus standard deviation, with significance reported for *p<0.05.

Tolerability of these treatments was explored in greater depth across an expanded scale of CFE. Mice received either procedural control (isoflurane anesthesia with no infusion) or IT-CM infusion of aCSF with varying tonicity (1X-9X) and were carefully monitored for ambulation time and gross tolerability. CFE treatments were very well tolerated at the 2X level and below, with a sub-2-minute ambulation time (**Figure 4A**). Mice that received 4X CFE were characterized by a mean ambulation time of 127 seconds, and we noted occasional respiratory depression that typically recovered within several breaths. As the aCSF tonicity was further increased, respiratory alteration became more frequent, and time to ambulation was much longer. We also observed abnormalities in meningeal vessels at the highest tonicities that likely reflect osmotic damage (not shown). A separate set of mice that received 1X, 2X, or 4X aCSF were subjected to MRI imaging prior to, immediately after, and 2 hrs after injection. Qualitative assessments based on 3D renderings of the ventricles yielded no change in shape of the ventricles for any infusion. Some quantitative differences in volume were observed as a function of aCSF concentration, although the changes were relatively minor (**Figure 4B**). The average baseline (i.e., pre-injection) ventricular volumes were measured to be 7.7 ± 0.15 µL, 8.6 ± 0.62 µL, and 8.9 ± 1.0 µL for the 1X, 2X, and 4X aCSF groups, respectively (**Figure 4C**). Immediately following aCSF administration, ventricular volume was observed to increase in all groups, with the 4X aCSF group showing the highest volume increase of 15.3 ± 12.0%. This is a statistically significant increase in ventricular volume, although it is below the threshold that would diagnose hydrocephalus in mice^18^. The change in ventricular volume was mitigated 2 hours post-injection, returning to within 10% of baseline value across all subjects tested. These data affirm that IT-CM infusion of 10uL of aCSF at a tonicity of up to 4X yields a transient, tolerable increase in ventricular volume that recovers within 2 hrs. Taken together, these results suggest that, for a 10uL IT-CM infusion, up to 4X aCSF is well tolerated by mice.

**Figure 4:**
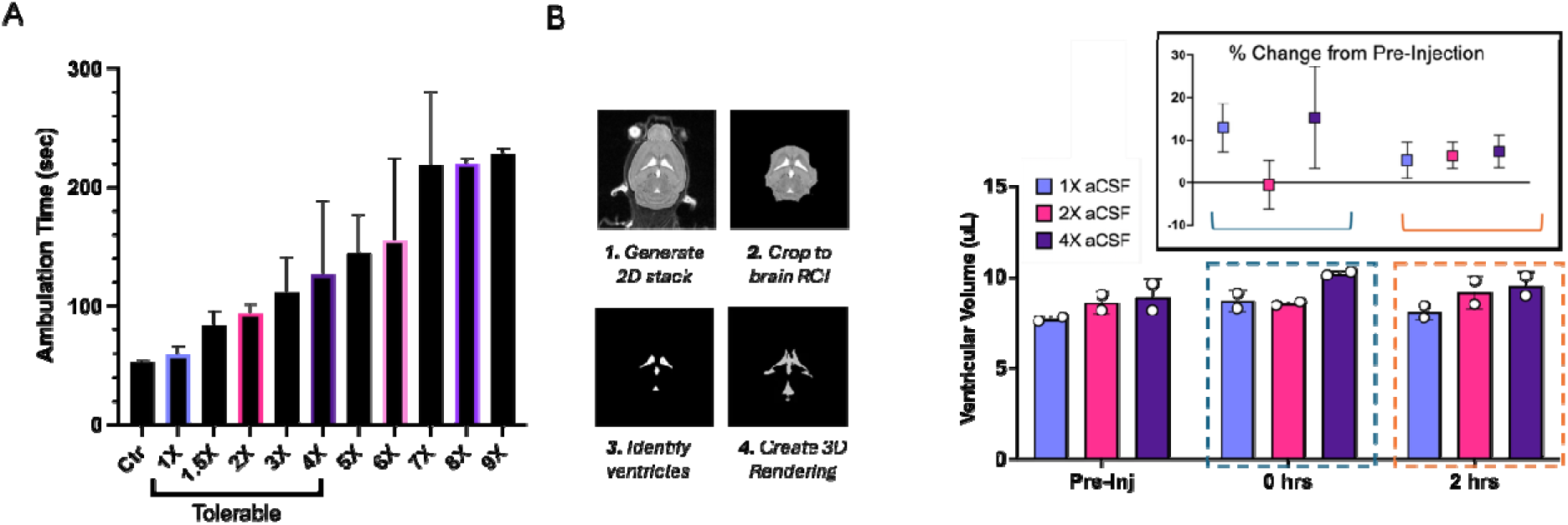
Application of CFE is tolerable and yields a mild, reversible increase in ventricular volume. Mice received procedural control (Ctr) or FNPs diluted in aCSF of varying tonicity (1X, 1.5X, 2X, 3X, 4X, 5X, 6X, 7X, 8X, 9X) as a 10uL infusion administered by the IT-CM route. (A) Up to 4X aCSF was considered tolerable, yielding a mean ambulation time of ∼2 minutes or less following brief isoflurane anesthesia. (B) MRI analyses demonstrated an increase in ventricular volume immediately after application of CFE that recovered within 2 hours of substance administration.

The next series of experiments sought to explore the kinetics of FNP distribution with whole-body widefield fluorescent imaging, which is a method in which any signal deep in tissue (for example, within the parenchyma or on the ventral aspect of the brain) will be highly attenuated relative to superficially detected signal. Mice received FNPs in 1X, 2X, or 4X aCSF under isoflurane anesthesia, after which they were immediately transferred to the imaging instrument. As the transfer introduces an unavoidable delay in imaging, the first measured time point is already at least 5 minutes after substance administration. Despite these limitations, IVIS imaging revealed several interesting observations. The dominant kinetics governing distribution were apparently fast, with the whole-head ROI peaking within 10 minutes of administration and the upper spinal cord slowly peaking between 1 and 2 hours after administration (see **Supplementary Figure 3**). This suggests initially fast distribution from the source compartment, with FNPs flushing through the subarachnoid space, settling into tissue, and clearing from dorsal surfaces of the brain soon after injection. FNPs appeared slowly in the upper spinal cord, with this ROI peaking at later time points, which suggests continued egress of FNPs from the cisterna magna.

Endpoint biodistribution of FNPs to peripheral organs was also assessed with IVIS (**Figure 5A**). The highest signal was measured in the kidney, liver, and lungs. Increasing dose of CFE tended to yield an increase in delivery to the kidneys and lungs, and there was a statistically significant decrease in delivery to the liver for the 4X condition compared to 1X. There was a very close relationship between delivery to liver versus lungs on a per-subject basis, but not between other organ pairs that were examined (**Supplementary Figure 4**).

**Figure 5:**
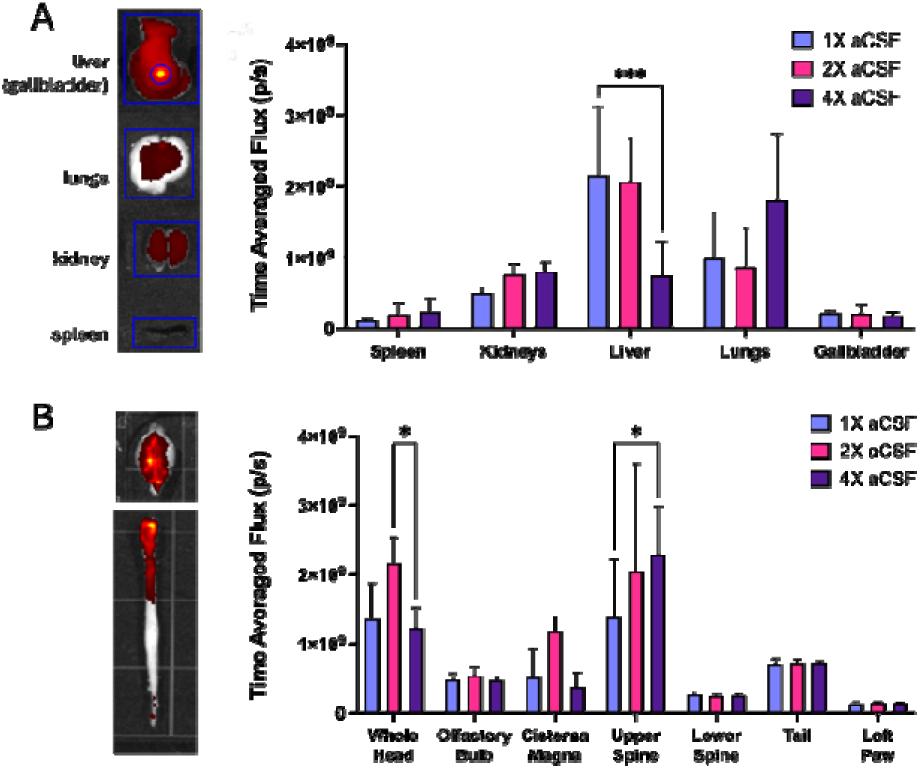
Whole organ fluorescence imaging reveals differences in peripheral and central tissue fate of FNPs following application of CFE. Mice received FNPs diluted in aCSF of varying tonicity (1X, 2X, or 4X) as a 10uL infusion administered by the IT-CM route. Time-averaged flux of intact tissue was assessed for both **(A)** peripheral and **(B)** central tissues. Data show mean plus/minus standard deviation (n=4 for 1X aCSF and n=3 each for 2X and 4X aCSF). Data were analyzed by two-way ANOVA with Dunnett’s multiple comparison test, and significance is reported for *p<0.05 and ***p<0.001.

Time-averaged flux measured in the whole-head ROI was significantly higher in the 2X aCSF group compared to the 1X and 4X aCSF groups (**Figure 5B**). This suggests that the 2X aCSF condition facilitates delivery of FNPs to the brain (or, understanding the limitations of whole-head imaging, to the ventral and easily detectable aspects of the brain) relative to the 1X and 4X conditions. When examining the time-averaged data of total flux within the head region over 2 hours, we observed significant differences in FNP delivery between the 2X and 4X groups (p<0.05) and nearly significant differences (p = 0.059) between the 2X and 1X aCSF groups. Additionally, higher total flux within the upper spine was observed for the 4X aCSF group when compared to the 1X and 2X aCSF groups (p<0.05). There were no significant differences in delivery to the lower spine, tail, and paw regions across CFE conditions.

The brain and spinal cord were next examined with fluorescent stereoscopy at higher magnification (**Figures 6** and **7**). Signal localization around vasculature was evident across all samples examined in the CNS, affirming our growing understanding that FNPs accumulate at PVS entry points into the parenchyma, i.e., Virchow-Robin channels. Across all samples imaged, vessel localization increased with increasing aCSF concentration. In the 1X aCSF group, FNP localization around the PVS appeared uniform and diffuse, and with increasing aCSF concentration, FNP distribution to the PVS became non-uniform, forming bright clusters proximal to the PVS. Within the brain (**Figure 6**), the olfactory bulb typically experiences high FNP accumulation, and increasing infusate tonicity yielded even higher FNP accumulation in this region. As previously observed, IT-CM administered FNPs preferentially localize to the ventral aspect of the brain, likely due to a high density of subarachnoid trabeculae and nerves running along the underside of the brain. IT-CM administered FNPs reach all CSF-exposed surfaces and localize especially with meningeal vessels. A relatively high concentration of FNPs were detected along the lateral brain vasculature, the Circle of Willis, basilar brainstem arteries, and posterior/anterior spinal arteries. Within the spinal cord (**Figure 7**), bright, isolated aggregates of fluorescence were observed in all subjects along the lateral edges and ventral aspects of the spine, with clear evidence of FNP movement into exiting nerve routes, another known egress pathway for CSF. Based on qualitative assessment, both vascular localization and nerve root localization were highest in the 2X group compared with the 1X or 4X conditions. Quantitatively, we observed an increase in medial-lateral distribution of signal across both the brain and the spinal cord for the 2X condition compared to the 1X condition (p<0.05) (**Figure 8**).

**Figure 6:**
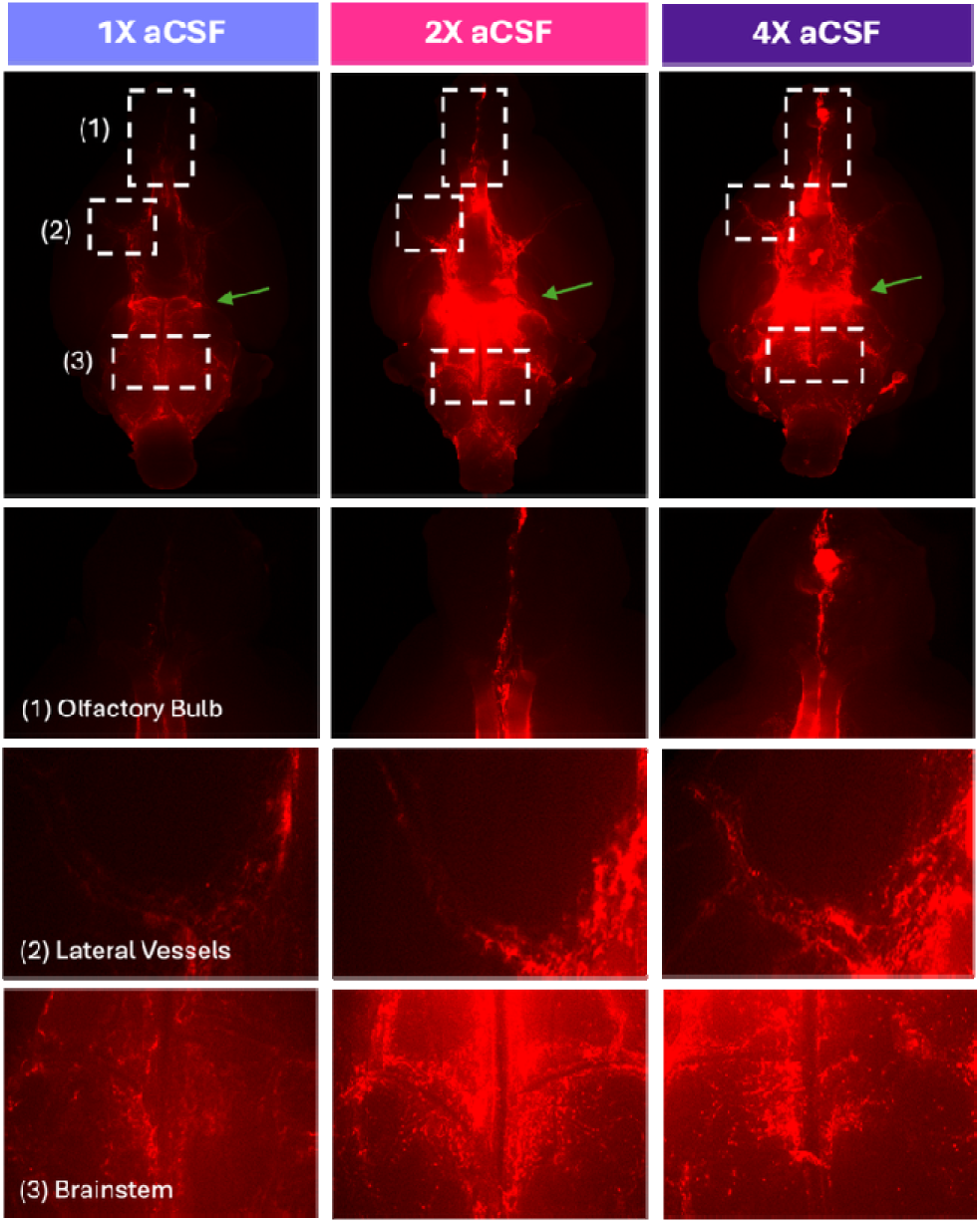
Following application of CFE, FNPs localize with ventral aspects of the brain. Regions of interest highlight consistently high delivery to the (1) olfactory bulbs, (2) lateral vessels of the Circle of Willis, and (3) brainstem. Images are representative of at least 3 experimental replicates

**Figure 7:**
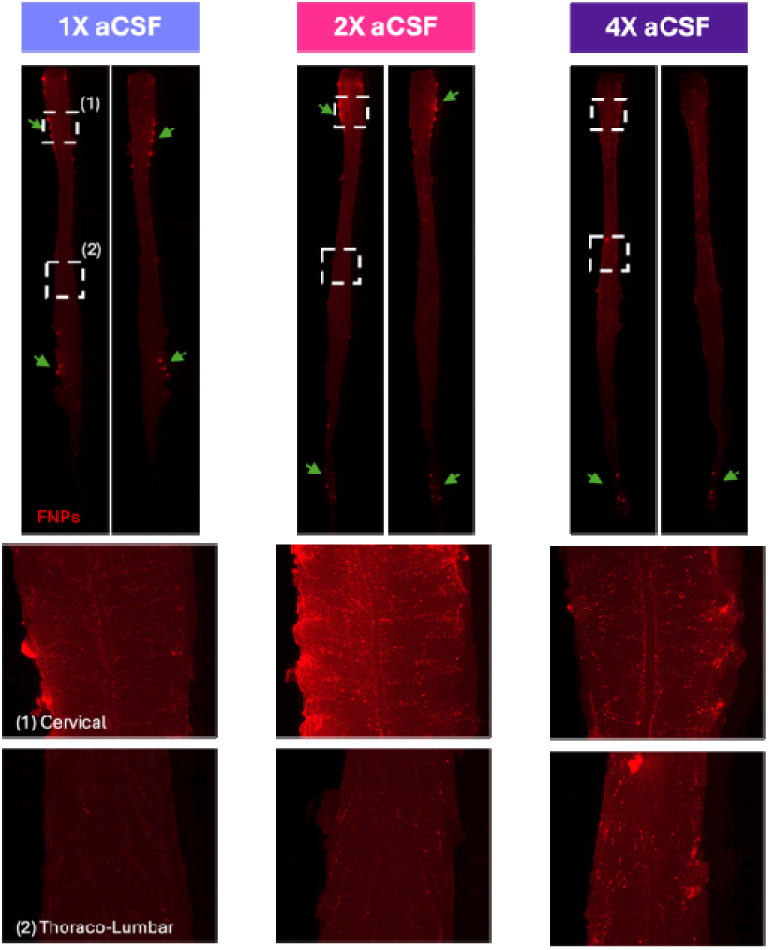
Following application of CFE, FNPs localize with upper portions of the spinal cord and exiting nerve roots. Signals are generally highest within the cervical region, which is adjacent to the IT-CM infusion site, and were lower in the thoraco-lumbar regions. Green arrows show examples of high localization of FNPs with exiting nerve roots. Images are representative of at least 3 experimental replicates.

**Figure 8:**
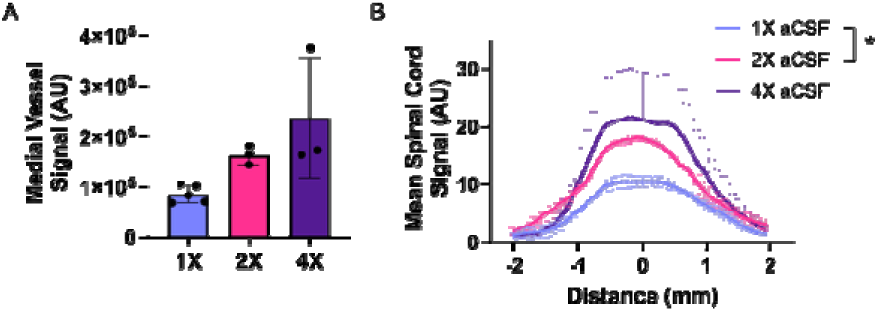
CFE tends to increase localization of FNPs with lateral vessels and improve their distribution in the spinal cord. This figure quantifies the complete data set represented in example images in Figures 6 (region (2)) and 7 (entire spinal cord)). (A) Shows whole ROI quantification of the medial vessels, representing the mean and standard deviation of n=3-5/group. (B) shows a medial-lateral line plot of FNP delivery to the entire spinal cord; the 2X condition yielded a significantly higher AUC compared to the 1X condition. These data show mean plus/minus standard error for n=3/group for visual clarity. Data were analyzed by one-tailed Mann-Whitney test, with significance reported for *p<0.05.

Our data suggest that increasing tonicity yields increased delivery of FNPs up to 4-6X aCSF and that delivery advantages are either lost or inconsistent as infusate tonicity is further increased. FNP signals became increasingly aggregated at the 6X level and above, which could be due either to selective localization within PVS entry points or aggregation within the SAS, yielding increased deposition on surfaces. We therefore examined the colloidal stability of FNPs maintained at 37^°^C in aCSF media of varying tonicity. The hydrodynamic diameter, zeta potential, and PDI of washed FNPs were measured by DLS (see **Supplementary Figure 5**). For the 1X and 1.5X aCSF groups, FNP hydrodynamic diameter remained relatively unchanged for up to 4 hours, with a variation of ≤ 10% from original values and low, stable PDI. We observed evidence of FNP aggregation when FNPs were maintained for longer periods of time in aCSF that was more concentrated than 1.5X. This aggregation was considered mild for the 2X aCSF group, with a ∼18nm increase by 1 hour and a ∼68nm increase by 2 hours. We note that our *in vivo* studies examined nanoparticle distribution for time points of only up to 2 hours after FNP administration. PDI trends were similar to changes in hydrodynamic diameter, with a minor increase in PDI up to 1 hour (PDI < 0.06 for the 2X group) and evidence for aggregation observed after 2 hours (PDI > 0.1 for the 2X group). In other words, for 1X-2X groups and considering time points of 2 hours or less, we consider these nanoparticles to be relatively stable colloids. Aggregation began to occur at the 4X level, with hydrodynamic diameters exceeding 1 µm and PDI exceeding 0.25. Taken in sum, these data confirm that 2X and below conditions are relatively stable, while 4X and above conditions may exhibit aggregation.

We examined cellular localization of FNPs as a function of aCSF tonicity on traditionally sliced tissues. FNP localization around vasculature was confirmed by spatial overlap of FNP fluorescence with CD31+ vascular staining (**Figure 9**). Consistent with the whole tissue imaging, application of CFE at the 2X or 4X levels altered the distribution of FNPs. Although no FNPs could be detected in parenchymal samples (at the exposure and gain settings used for this experiment) for the 1X control, FNPs were observed to penetrate the parenchyma in the 2X condition, and to achieve deep penetration into the PVS in the 4X condition, traveling up to 300µm from the meningeal border / subarachnoid space.

**Figure 9.**
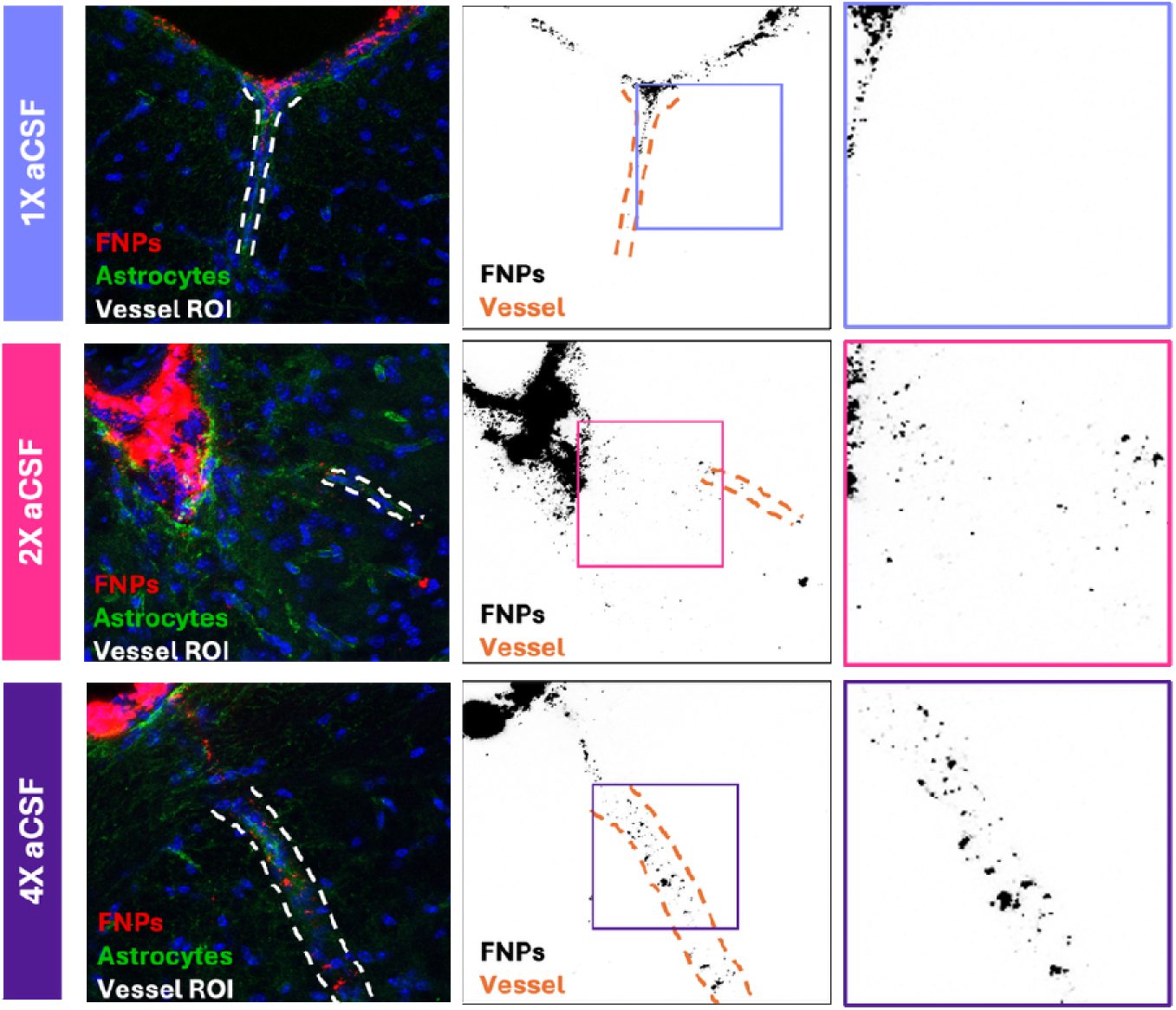
Application of CFE alters the parenchymal fate of FNPs after IT-CM administration. Sliced tissues were stained for DAPI (blue) and AQP4 (green). Vessels, outlined with dashed white lines, were identified by the characteristic shape of AQP4+ astrocytic end feet; the red channel is inverted in black and white with zoomed inset to highlight delivery features. Under these imaging conditions, no FNPs were detected in the PVS or parenchyma of mice that received a standard 1X aCSF infusion. Following application of CFE, FNPs were observed to enter the brain parenchyma (2X) and to transit hundreds of microns into the PVS (4X).

In the last series of experiments, we queried whether the improvements in distribution observed for 100nm polystyrene nanoparticles would hold for differently sized colloids. FNPs of variable hydrodynamic diameter (20, 40, or 100nm) were infused into healthy mice by the IT-CM route (**Figure 10**). When infused under standard conditions, in 1x aCSF, FNPs were observed at a high concentration on the periphery of the brain, in tissue directly adjacent to the subarachnoid space, ventricles, or cisterns. When infused under CFE conditions, in 2x aCSF, the shape of the cerebral ventricles was often well delineated, with qualitative assessments confirming that the improvements in CNS delivery robustly observed for 100nm FNPs are generalizable to colloids of other sizes. The benefits of CFE may be higher for 40nm FNPs compared to 100nm FNPs, although answering this question with quantitative rigor remains the subject of future work.

**Figure 10.**
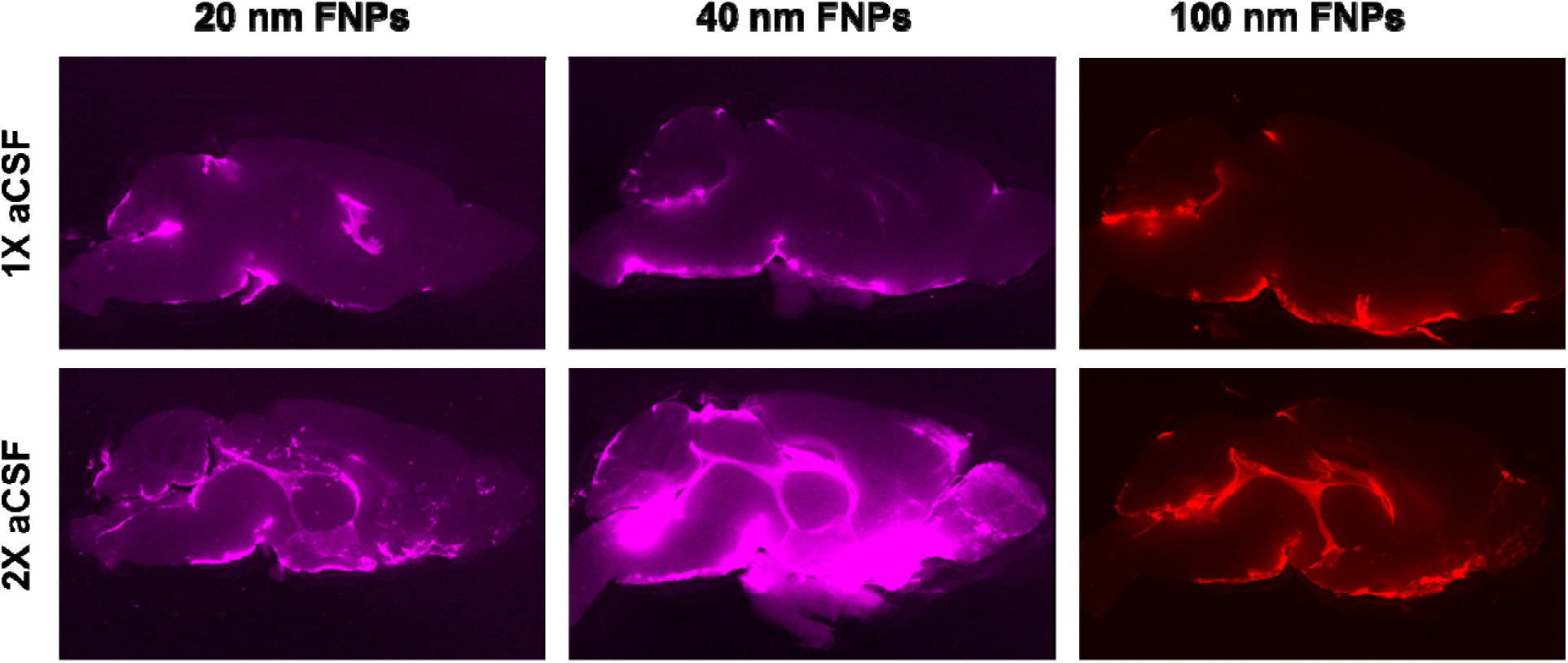
CFE yields generalizable improvements in brain delivery of variably sized FNPs. Mice received an IT-CM injection of 20, 40, or 100nm FNPs, after which brains were extracted and sliced into thick sagittal sections. CFE yielded qualitative improvements in penetration of FNPs through the ventricular system and into deep tissue regions.

## Discussion

We sought to manipulate the tonicity of intrathecal infusion media for the purpose of improving CNS drug delivery. Our data provide evidence to support the overarching hypothesis that intrathecal infusion of hypertonic fluid can enhance colloid delivery to specific tissues of the CNS. Our data establish a dose-response relationship between aCSF concentration and various aspects of nanoparticle localization in the brain and spinal cord, demonstrating that infusion of up to 4X aCSF is tolerable and can provide significant benefits for CNS drug delivery. In totality, we describe our approach as part of a family of methods termed CSF Flow Enhancement, or CFE, a growing field of study regarding the manipulation of CSF production, flow, and clearance for therapeutic purposes. The following sections discuss these novel findings, describe limitations of the current work, and speculate on the mechanisms by which hypertonic aCSF achieves CFE.

We posit that infusion of hypertonic fluid into the cisterna magna, which is immediately adjacent to the CP of the 4^th^ ventricle, stimulates a flush of fluid production that washes nanoparticles across and into CNS tissues. An alternative hypothesis is that fluid is drawn out of tissue, but our data do not support this expectation: (1) FNPs administered in 1X aCSF do not enter the parenchyma, whereas FNPs do when administered in 2X and 4X aCSF do (interstitial fluid egress would result in tightened extracellular spaces, limiting rather than promoting delivery); and (2) Entry of FNPs into the cervical lymphatic system supports glymphatic-mediated transport, which would be disrupted if tissues were dehydrated.

Our data establish that the composition of the aCSF medium used to administer nanoparticles by the IT route plays a significant role in determining nanoparticle fate. The selection of an infusion medium is in itself a complex consideration, with wide variation in clinical and preclinical approaches spanning the use of distilled water, PBS, Ringer’s, or variably composed aCSF. Although our recipe for aCSF was derived from established aCSF formulations, none of these formulations were created to exactly mimic the CSF composition of mice or humans^13–16^, having been developed instead for the purpose of maintaining tissue viability in brain slice experiments. The formulations utilized here are therefore not expected to bear properties identical to native CSF but instead provide a first-pass approach for adjusting total infusion medium tonicity by scaling all solutes equivalently; it remains the subject of future work to selectively manipulate individual ions. We prepared aCSF with an average pH value of ∼5.11 ± 0.19 across all aCSF concentrations (see **Supplementary Figure 5**). Although this is considered an acidic pH, the infusions were well tolerated by mice through the 4X level. Given that the chemosensory capability of CP includes both osmolality and pH, it remains the subject of future work to untangle the independent effects of tonicity versus acidity. MRI findings show that infusion of hypertonic aCSF transiently increases the volume of the cerebral ventricles but does not produce permanent changes. A previous study defined mice with a ventricular volume of greater than 14mm^3^ (i.e., 14µL) as having a hydrocephalus condition or, more conservatively, an enlarged ventricular volume^18^. Since our mice are the same strain, sex, and age, we are confident using this report as a reference point to establish the safety of the CFE method.

DLS data demonstrated a relationship between increasing aCSF tonicity and NP stability, which we have previously reported^3^. Increasing solution osmolality can lead to NP aggregation through the adsorption of oppositely charged ions onto the NP surface, which neutralizes surface charge and enables a higher rate of nanoparticle-nanoparticle contact, leading to aggregation^19^. The FNPs used in this study were carboxylated, and so they bear an overall negative charge that would attract positively charged ions present in aCSF, including Na^+^, K^+^, Ca^2+^, and Mg^2+^. Aggregation was consistently observed at higher osmolality conditions (i.e., 4X, 6X, and 8X aCSF concentrations) across all time points, but it was not observed for the 2X or lower condition. The question of whether aggregation contributes to the enhanced FNP delivery seen for CFE conditions is important: we posit that FNP aggregation would serve to *reduce* not *enhance* the spatial distribution of colloids as they navigate through tortuous networks of collagen-rich, trabecular fibers that infiltrate the subarachnoid space. Appearance of FNPs in distal regions of the CNS and within deep parenchymal targets emphasize that FNPs remain small enough and mobile enough to be transported across and into tissues for the 4X condition. However, we did notice declining benefit of CFE in the 6X and 8X groups; given the appearance of particularly large puncti in some of these higher tonicity samples (data not shown), we posit that aggregation becomes a problem at the 6X or 8X level, presumably due to greater resistance to bulk flow, thereby leading to restricted movement and heterogeneous distribution. Whether aggregation of FNPs at higher tonicity contributes to lengthy ambulation times at these higher levels of CFE remains to be determined.

Through IVIS and *ex vivo* fluorescence, we noted that application of CFE yielded an increased delivery of FNPs throughout the CNS and into peripheral tissues, including CLNs. Additionally, there was a tendency for increased delivery to the kidneys and lungs upon application of CFE. We also noted major differences in the tissue-level fate of FNPs after application of CFE, including the appearance of FNPs within the parenchyma (2X aCSF) and PVS of the parenchyma (4X aCSF), which we have never observed under standard (1X aCSF) conditions. In addition, the 4X aCSF group exhibited higher total flux within the upper spinal cord in comparison to other aCSF groups from IVIS imaging, suggesting a shift in bulk flow towards the spine. Considering sagittal views of the CNS, we also observed delineation of the cerebral ventricles in the 2X and 4X groups, only, not the 1X control circumstance. In totality, these data suggest that application of CFE facilitates fluid production and egress from the CNS through the glymphatic and lymphatic systems.

We are not the only research group interested in manipulating CNS fluids; in fact, a variety of CFE methods exist that operate through pharmacological, biophysical, or physical means (examples provided in **Table 1**). Increasingly strong evidence supports the expectation that CFE will be an important technique technique: in one example, inhibition of motile cilia with intrathecal lidocaine enabled 40-fold higher concentration of temozolomide delivery to murine brain tumor, by stagnating perivascular clearance of the chemotherapeutic^20–23^. Although we have primarily studied the use of CFE to drive exogenous substances *into* the parenchyma, a corresponding technique, i.e., using CFE to drive endogenous substances *out of* the parenchyma, may be equally significant. This is of particular interest given growing field knowledge that damage to the glymphatic system, triggered either by brain injury or by neurodegenerative processes, leads to impaired waste clearance and buildup of substances that contribute to neuroinflammation and neuronal death. Thus, CFE is emerging as a powerful approach by which this toxic buildup can be cleared for neuroprotective purposes.

**Table 1:** Published approaches for achieving CFE in murine models.

| Approach | Examined in | Observations |
| --- | --- | --- |
| IV Hypertonic saline | Healthy<sup>24</sup> | Reduced intracranial pressure; clearance of tracers from PVS |
| IV Pan-adrenergic inhibition | Ischemic stroke<sup>25</sup>; Epilepsy<sup>26</sup>; TBI<sup>27</sup> | Improved outcome; decreased cell debris; clearance of p-tau |
| IV Ethanol | Addiction<sup>28</sup> | J-shaped dose response; clearance of tracers from PVS |
| IV SalB/Rg1 | Ischemic Stroke<sup>29</sup> | Normalized AQP4; enhanced solute transport; reduced edema |
| IT Lidocaine | Brain cancer<sup>21</sup> | Inhibition of motile cilia; 40x increase in temozolomide delivery |
| IT Mannitol | Healthy<sup>20</sup> | Enhanced PVS penetration of antibodies |
| IP AQP4 inhibition | Healthy<sup>30</sup> | Brain-region specific redirection of PVS fluid |
| Multisensory stimulation | Alzheimer's<sup>31</sup> | Increased AQP4 polarization; increased amyloid beta clearance |
| Physical movement | Healthy<sup>22</sup> | Voluntary running (mouse); enhanced tracer distribution in fixed tissue |
| Focused ultrasound | Healthy<sup>23</sup> | Enhanced transport of tracer along PVS; entry into the parenchyma |

In conclusion, we present an investigation into the use of intrathecal, hypertonic aCSF for enhancing CSF transport of intrathecally administered nanoparticles. Our results demonstrate that the distribution of nanoparticles in the CNS depends on the tonicity of the infusion medium, with 2X and 4X conditions yielding increased association of FNPs with the surfaces, perivascular spaces, and bulk tissues of the brain and spinal cord. Taken together, these data demonstrate that manipulation of infusion media tonicity can be a powerful tool for driving nanoparticle delivery to specific tissue regions of the CNS.

## Acknowledgements

We gratefully acknowledge funding for this work from Ian’s Friends Foundation as well as the National Institute for Neurological Disease and Stroke (NINDS), the Eunice Kennedy Shriver National Institute of Child Health and Disease (NICHD), and the National Institute on Aging (NIA) at the National Institutes of Health (R01NS111292, R01NS116657, R01HD099543, R21AG089301).

