## Supplemental Data for "Intrathecal infusion of hypertonic fluid enables CSF Flow Enhancement (CFE) to facilitate nanoparticle delivery to the brain and spinal cord"

### Slide 1
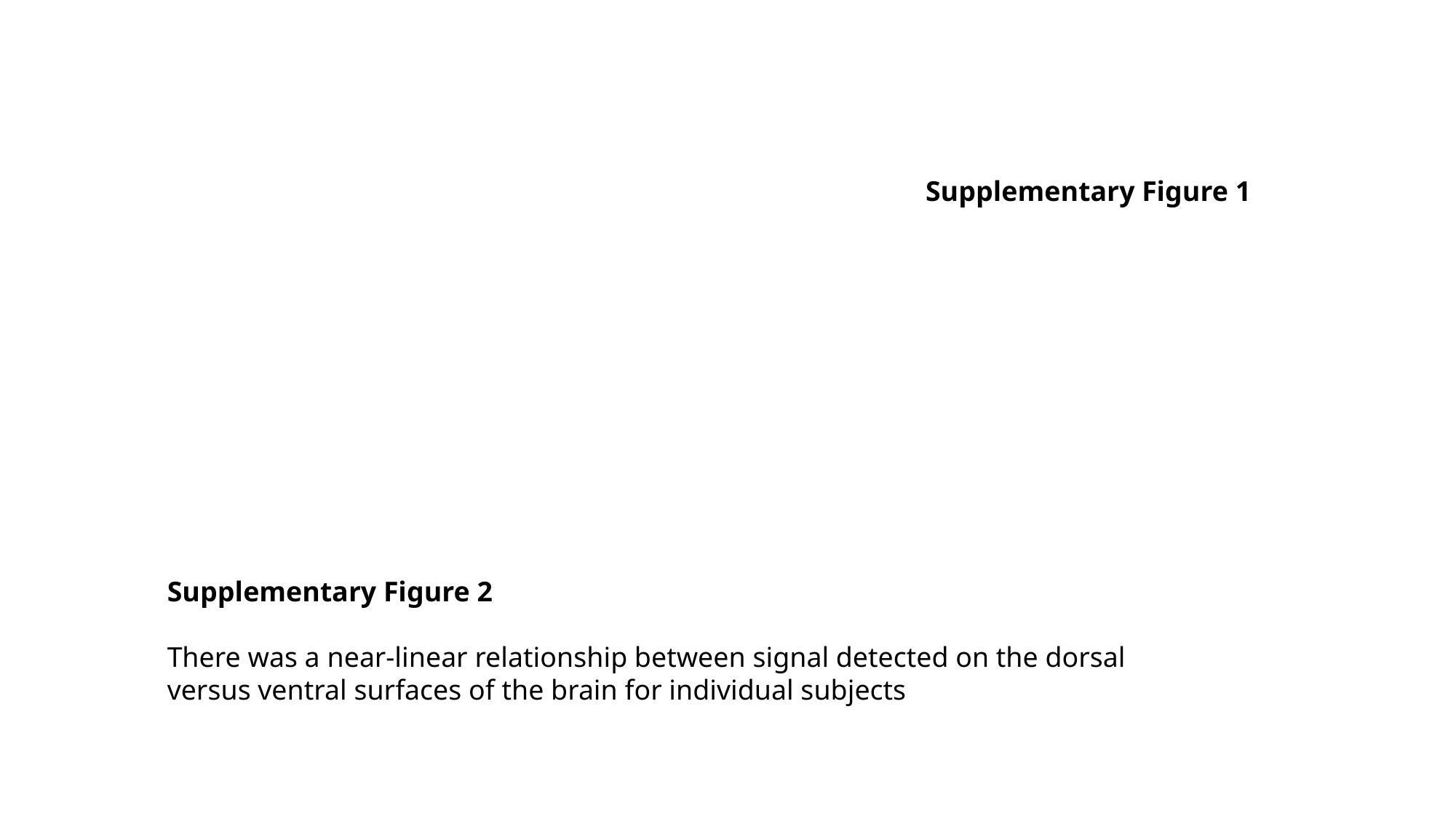

Supplementary Figure 1
Supplementary Figure 2
There was a near-linear relationship between signal detected on the dorsal versus ventral surfaces of the brain for individual subjects

### Slide 2
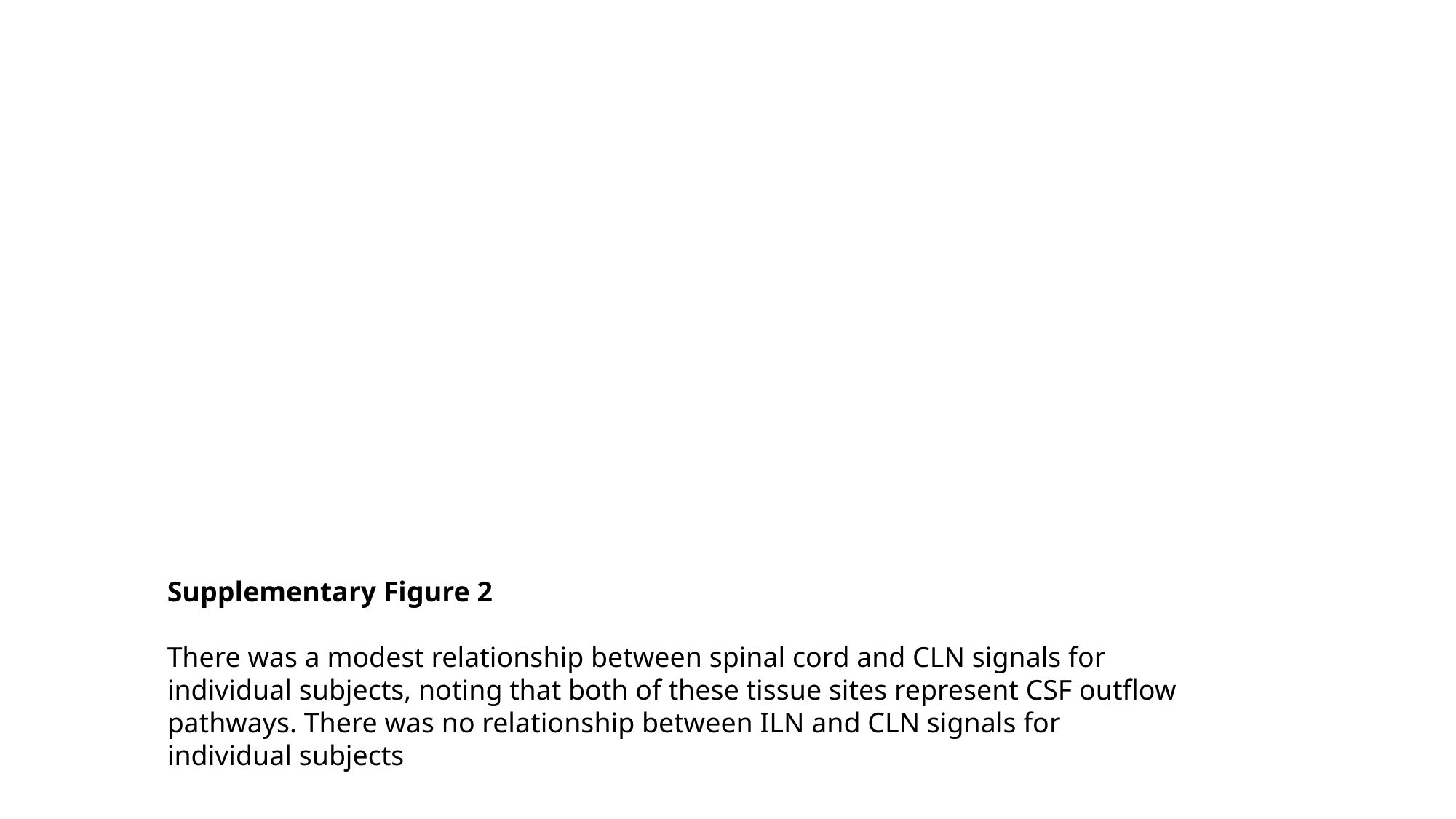

Supplementary Figure 2
There was a modest relationship between spinal cord and CLN signals for individual subjects, noting that both of these tissue sites represent CSF outflow pathways. There was no relationship between ILN and CLN signals for individual subjects

### Slide 3
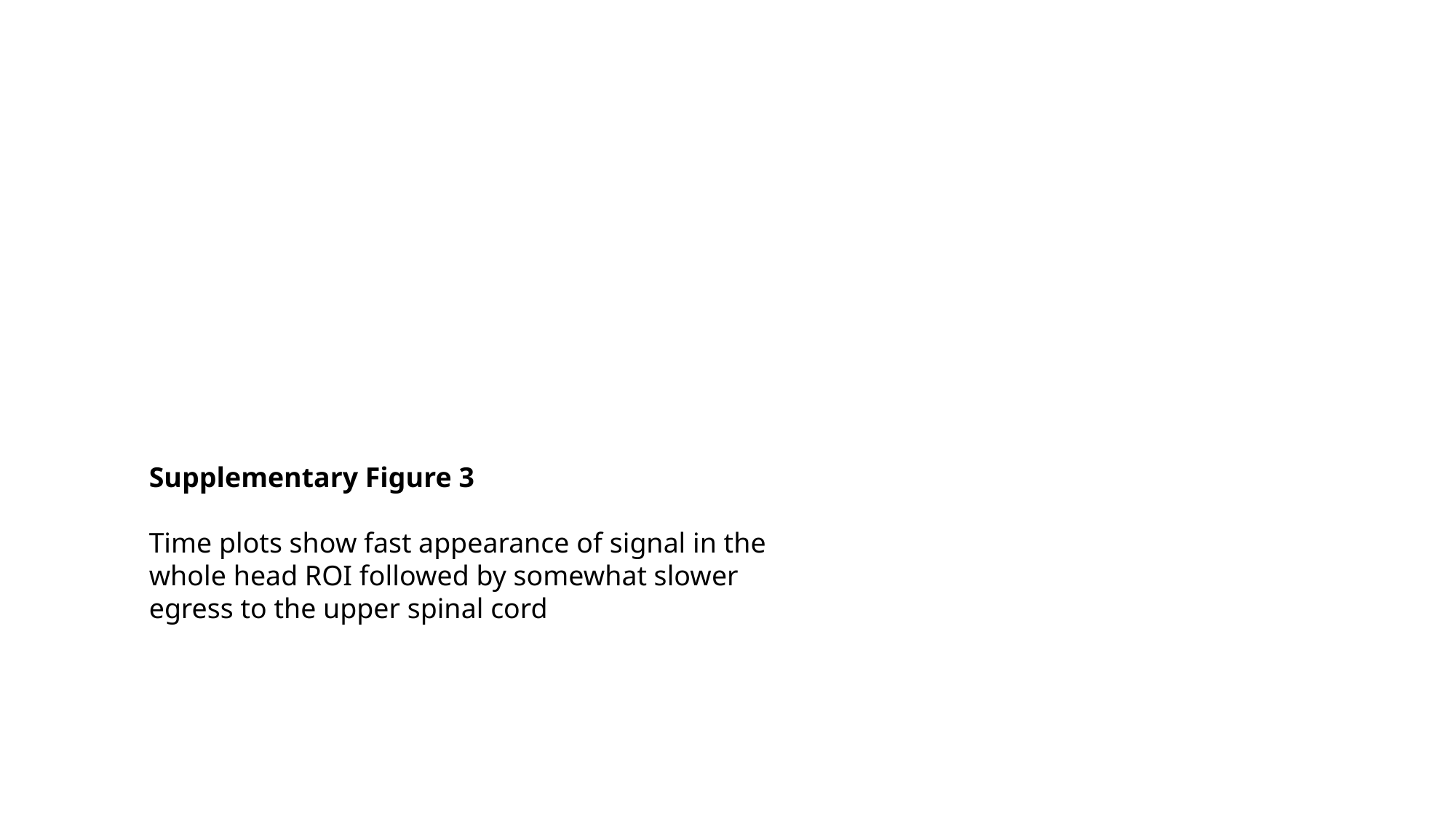

Supplementary Figure 3
Time plots show fast appearance of signal in the whole head ROI followed by somewhat slower egress to the upper spinal cord

### Slide 4
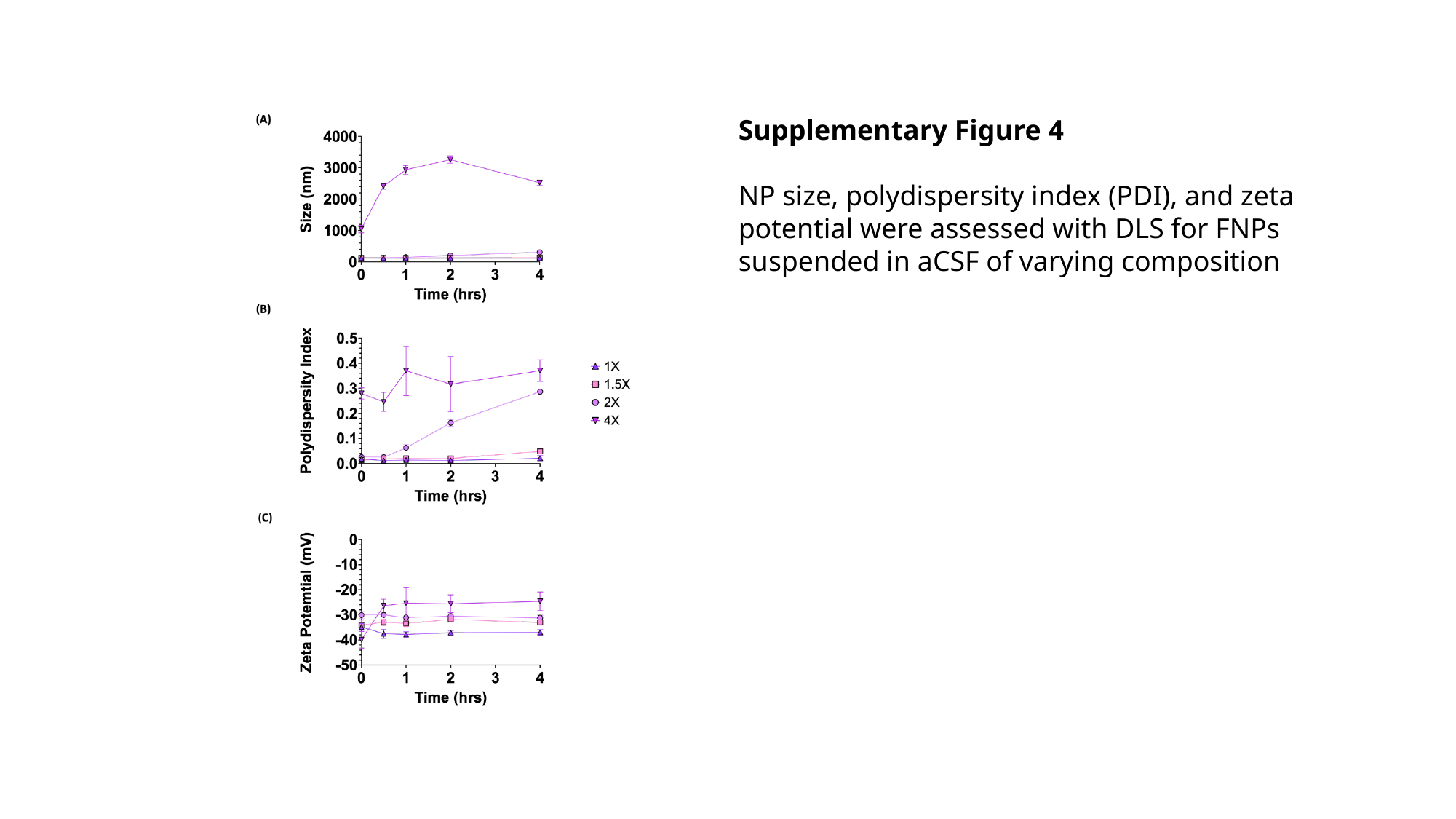

Supplementary Figure 4
NP size, polydispersity index (PDI), and zeta potential were assessed with DLS for FNPs suspended in aCSF of varying composition

### Slide 5
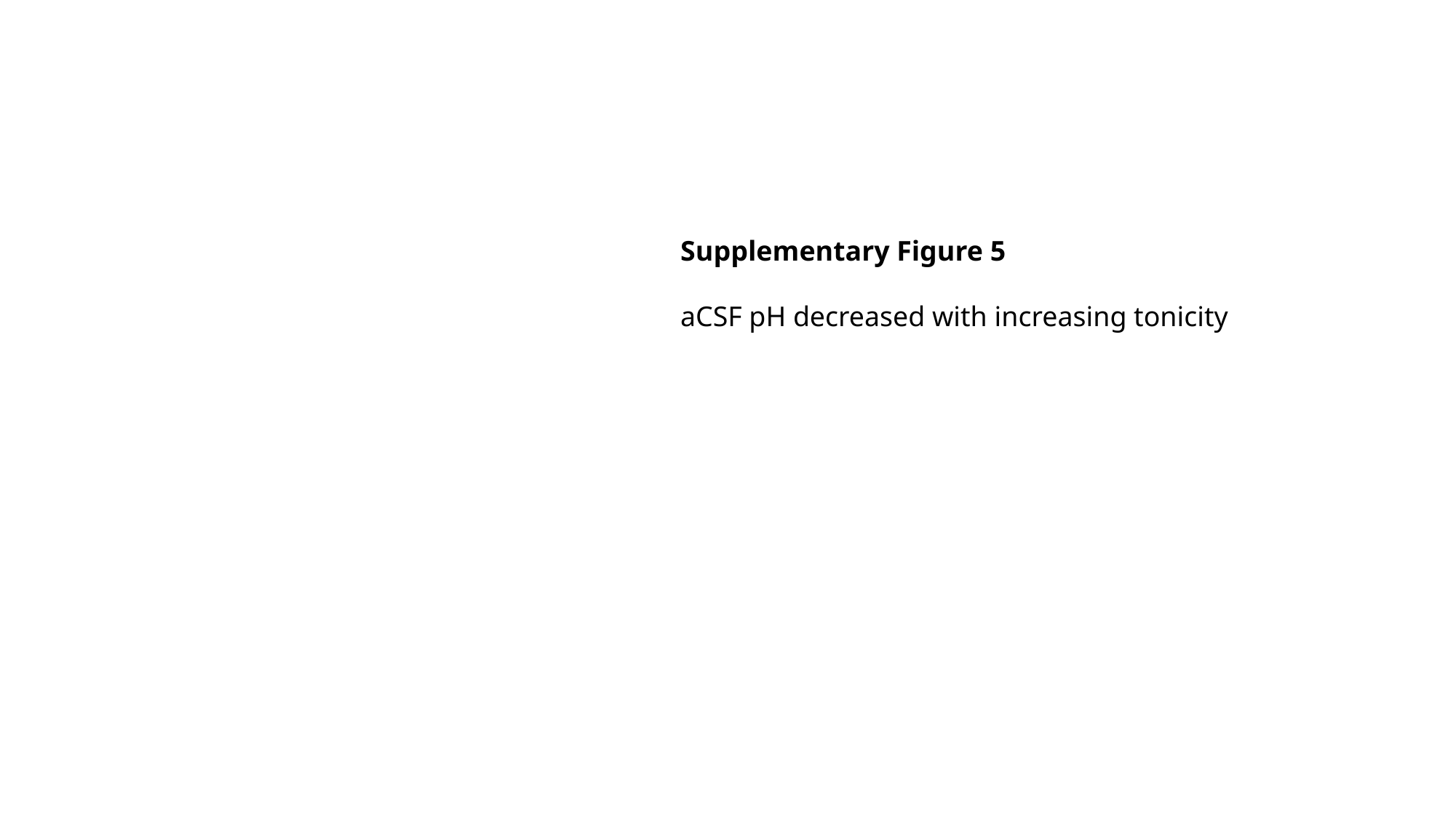

Supplementary Figure 5
aCSF pH decreased with increasing tonicity
